# Short N-terminal disordered regions and the proline-rich domain are major regulators of phase transitions for full-length UBQLN1, UBQLN2 and UBQLN4

**DOI:** 10.1101/2023.09.27.559790

**Authors:** Thuy P. Dao, Anitha Rajendran, Sarasi K. K. Galagedera, William Haws, Carlos A. Castañeda

**Affiliations:** Departments of Biology and Chemistry, Syracuse University, Syracuse, NY 13244, USA; Interdisciplinary Neuroscience Program, Syracuse University, Syracuse, NY 13244, USA; BioInspired Institute, Syracuse University, Syracuse, NY 13244, USA

**Keywords:** Phase separation, UBQLN1, UBQLN2, UBQLN4, N-terminal disordered region, proline-rich region, charge state, epitope tags

## Abstract

Highly homologous ubiquitin-binding shuttle proteins UBQLN1, UBQLN2 and UBQLN4 differ in both their specific protein quality control functions and their propensities to localize to stress-induced condensates, cellular aggregates and aggresomes. We previously showed that UBQLN2 phase separates *in vitro*, and that the phase separation propensities of UBQLN2 deletion constructs correlate with their ability to form condensates in cells. Here, we demonstrated that full-length UBQLN1, UBQLN2 and UBQLN4 exhibit distinct phase behaviors *in vitro*. Strikingly, UBQLN4 phase separates at a much lower saturation concentration than UBQLN1. However, neither UBQLN1 nor UBQLN4 phase separates with a strong temperature dependence, unlike UBQLN2. We determined that the temperature-dependent phase behavior of UBQLN2 stems from its unique proline-rich (Pxx) region, which is absent in the other UBQLNs. We found that the short N-terminal disordered regions of UBQLN1, UBQLN2 and UBQLN4 inhibit UBQLN phase separation via electrostatics interactions. Charge variants of the N-terminal regions exhibit altered phase behaviors. Consistent with the sensitivity of UBQLN phase separation to the composition of the N-terminal regions, epitope tags placed on the N-termini of the UBQLNs tune phase separation. Overall, our *in vitro* results have important implications for studies of UBQLNs in cells, including the identification of phase separation as a potential mechanism to distinguish the cellular roles of UBQLNs, and the need to apply caution when using epitope tags to prevent experimental artifacts.

**Highlights:** 1) Full-length UBQLN1, UBQLN2 and UBQLN4 all exhibit LCST phase transitions but to different degrees.

2) The N-terminal region (N-terminal to the UBL) substantially regulates UBQLN phase separation.

3) Removal of the disease-associated proline-rich (Pxx) region in UBQLN2 removes the strong temperature dependence of UBQLN2 phase separation.

## Introduction

UBQLNs are ubiquitin-binding shuttle proteins critical to mediating protein quality control in cells (Zheng et al., 2020; Zientara-Rytter and Subramani, 2019). Highly homologous UBQLN1, UBQLN2 and UBQLN4, the predominant forms of UBQLNs in human cells, are associated with several neurodegenerative diseases. Notably, several mutations in UBQLN2 cause dominant X-linked form of amyotrophic lateral sclerosis (ALS) and frontotemporal dementia (FTD) (Daoud et al., 2012; Deng et al., 2011; Fahed et al., 2014; Synofzik et al., 2012; Teyssou et al., 2017). Moreover, a variant of UBQLN4 is linked to ALS, and dysregulation of UBQLN1 has been implicated in Alzheimer’s disease (Adegoke et al., 2017; Edens et al., 2017; Mah et al., 2000; Zhang et al., 2022). UBQLNs are implicated in Huntington’s and Parkinson’s diseases (Gerson et al., 2020; Mori et al., 2012; Wang et al., 2006). In all of these cases, UBQLNs were found in protein-containing inclusions that suggest dysregulated protein quality control as a contributing mechanism to disease. Therefore, it is important to discern the biophysical properties and molecular mechanisms underlying how UBQLNs function in cells.

We and others recently determined that UBQLN2 colocalizes with biomolecular condensates called stress granules, independently forms stress-induced condensates in cells, and undergoes phase separation *in vitro* at physiological concentrations (Alexander et al., 2018; Dao et al., 2018; Gerson et al., 2021; Riley et al., 2021; Sharkey et al., 2018). We further demonstrated that the STI1-II region (STI3/4) of UBQLN2 is central to its ability to oligomerize, phase separate and form stress-induced condensates. The other UBQLN2 paralogs, UBQLN1 and UBQLN4, exhibit similar length and domain architecture, including the highly conserved oligomerization STI1-II domain (Fig. 1A). Like UBQLN2, the C-terminal STI1-II-containing regions of UBQLN1 and UBQLN4 are capable of phase separation *in vitro* (Gerson et al., 2021). UBQLN1 and UBQLN4 also form condensates and/or aggregates in cells, but to different extents (Gerson et al., 2021; Heir et al., 2006; Mah et al., 2000; Regan-Klapisz et al., 2005). As dysregulation of condensates is thought to lead to protein inclusions in age-related diseases (Alberti and Hyman, 2021; Elbaum-Garfinkle, 2019), we surmise that the different abilities of UBQLN1, UBQLN2 and UBQLN4 to phase separate and form condensates contribute to and differentiate their cellular functions and disease states.

**Figure 1.**
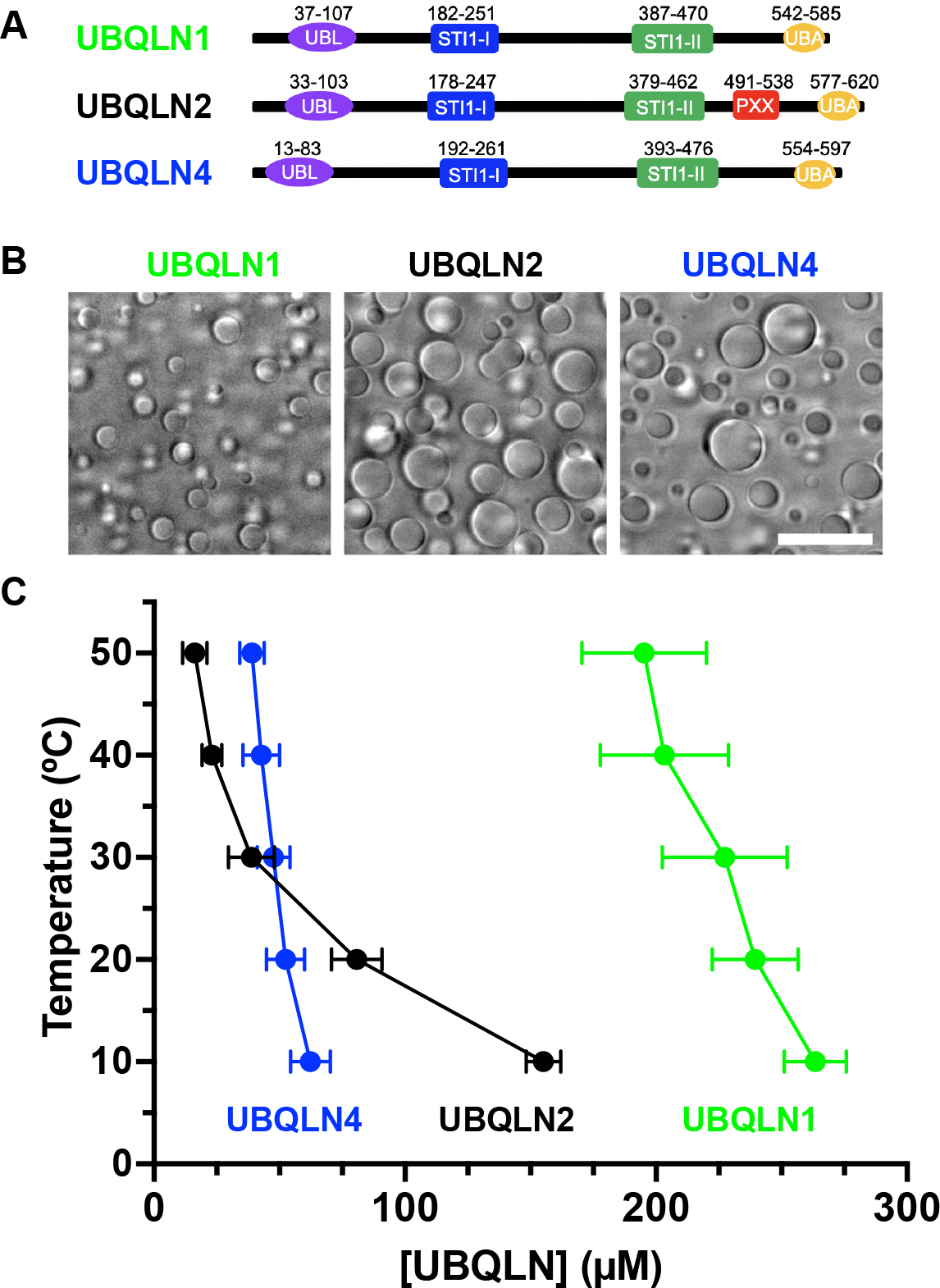
UBQLN1, UBQLN2 and UBQLN4 exhibit different phase behaviors. (A) Domain architectures of UBQLN1, UBQLN2 and UBQLN4. The proline-rich repeat (Pxx) region is unique to UBQLN2. (B) Brightfield microscopy shows 200 µM solutions of the three proteins in buffer containing 20 mM NaPhosphate, 200 mM NaCl, 0.5 mM TCEP, and EDTA (pH 6.8) after incubation at 30 °C for 5 minutes. Imaging was done at the coverslip. Scale bar, 10 µm. (C) Phase diagrams determined from c_sat_ measurements as a function of temperature. Phase separation is observed above these coexistence lines. Error bars represent SD over at least four trials from two different proteins preps.

In this study, we set out to quantitatively compare the phase transitions of full-length UBQLN1, UBQLN2, and UBQLN4 as well as determine the sequence features that distinguish them. Here, we expressed and purified tagless UBQLNs from bacteria and obtained temperature-concentration phase diagrams, which showed distinct phase transitions for the three proteins. Using deletion constructs and variants, we determined the UBQLN regions that contribute to the differences in the phase transitions of UBQLN1, UBQLN2 and UBQLN4. In addition, we demonstrated that certain epitope tags placed on the N-termini of UBQLNs substantially alter the phase transitions of the wild-type proteins. Results from previous studies on UBQLNs in cultured cells could be complicated by overexpression and location of epitope or fluorescent tags on UBQLNs that are used for visualization. Both of these issues can affect condensate formation, material properties, and subcellular localization (Alberti et al., 2019; Riley et al., 2021; Uebel and Phillips, 2019). Moreover, due to high sequence identities (e.g., 74% for UBQLN1/UBQLN2, and 56% for UBQLN2/UBQLN4) among the three UBQLNs, most antibodies are not specific for just one UBQLN which makes detection of endogenously expressed proteins ambiguous (Lin et al., 2021; McDowell et al., 2023). The results from this study will enable better experimental designs for future functional studies of UBQLNs in cells and provide a framework to examine the effects of epitopes on other phase-separating systems.

## Results

### UBQLN1, UBQLN2 and UBQLN4 exhibit different phase behaviors

We recently showed that full-length UBQLN2 undergoes phase separation under physiological conditions. Moreover, C-terminal constructs of all three UBQLNs undergo phase separation at high salt concentrations (Gerson et al., 2021). Therefore, we hypothesized that full-length UBQLN1 and UBQLN4 also undergo phase separation. To systematically study the three proteins, we expressed and purified tagless full-length UBQLN1, UBQLN2 and UBQLN4 using our salting out method (see Materials and Methods) (Fig. S1). SEC-MALS-SAXS (size-exclusion chromatography coupled with multiangle light scattering and small-angle X-ray scattering) data show that all three proteins exist as dimers and have similar conformations in solution, as revealed by R_g_ (radius of gyration) and P(r) measurements (Figs. S2-4). The ability to self-associate (via the STI1-II domain) is necessary for UBQLN2 to phase separate (Dao et al., 2018). We examined the phase separation of UBQLNs by adding salt to purified protein solutions. All three proteins formed liquid-like droplets in 200 mM NaCl, but to different extents (Fig. 1B). Specifically, UBQLN2 and UBQLN4 droplets were larger than those of UBQLN1, suggesting a higher phase separation propensity for UBQLN2 and UBQLN4.

To directly and quantitatively compare the phase behaviors of UBQLN1, UBQLN2 and UBQLN4, we obtained temperature-concentration phase diagrams for the three proteins by measuring saturation concentrations (c_sat_) at different temperatures (Fig. 1C). Consistent with our microscopy data, UBQLN2 and UBQLN4 phase separated much more readily (at lower c_sat_) than UBQLN1. These data are consistent with previous observations for UBQLN1 compared to UBQLN2/4 *in vitro* and in cells (Buel et al., 2023; Gerson et al., 2021). Moreover, the c_sat_ for all three proteins decreased with increasing temperatures, consistent with lower critical solution temperature (LCST) phase behavior. However, UBQLN2 phase separation exhibited a greater temperature dependence than UBQLN1 and UBQLN4 (over the temperature range examined), especially at lower temperatures between 10 °C and 30 °C. These differences in the phase behaviors of the three UBQLNs were surprising to us. Since UBQLN1 and UBQLN2 exhibit 74% sequence identity whereas UBQLN2 and UBQLN4 exhibit 56% sequence identity, we had expected UBQLN1 and UBQLN2 to have more similar phase behaviors. Therefore, we surmise that the differences within individual domains of the UBQLNs distinguish how each protein phase separates.

### UBQLN2 Pxx region and N-terminal disordered regions distinguish UBQLN phase transitions

Through domain comparisons and the specificity of UBQLN2 antibodies, we identified two distinguishing regions within the three UBQLNs. First, UBQLN2 is well-known to differ from its paralogs by the presence of a unique Pxx (proline-rich repeat) region, where most disease-linked mutations are harbored (Deng et al., 2011; Zheng et al., 2020). Secondly, the three UBQLNs also contain distinguishable disordered N-termini, as determined by the specificities of UBQLN antibodies. Western blots using commercially available UBQLN2 antibodies show many with cross reactivity to UBQLN1, which migrates slightly more quickly than UBQLN2 on a SDS-PAGE gel (McDowell et al., 2023). We tested three UBQLN2-specific antibodies, CST#85509, 23449-1-AP and NBP2-25164, that do not recognize UBQLN1 and UBQLN4 (Fig. S5). Using several UBQLN2 deletion constructs, we determined that these antibodies primarily recognize epitopes within UBQLN2 residues 1-33, an intrinsically disordered region that is N-terminal to the UBL domain. Removal of these residues (e.g., UBQLN2 33-624) renders these three antibodies unable to recognize UBQLN2, thus indicative of the uniqueness of the N-terminal region of UBQLNs. We note that this region has been understudied and has been largely considered part of the UBL domain that is highly homologous across the three UBQLNs. Therefore, we focused on how the Pxx and N-terminal disordered regions contribute to regulating the phase behaviors of the UBQLNs.

Removal of the Pxx region from UBQLN2 (UBQLN2-Pxx) abolished the sharp temperature dependence observed for full-length UBQLN2 phase diagram at low temperatures (Fig. 2A). Surprisingly, the phase diagram for UBQLN2-Pxx was nearly identical to that of UBQLN4, which exhibits a low temperature dependence in its phase behavior between 10 and 50 °C. Conversely, adding a large portion of UBQLN2 Pxx into UBQLN4 (UBQLN4+Pxx, Fig. S6) modestly increased the temperature dependence of UBQLN4 phase behavior (Fig. 4A). Together, these data demonstrate the importance of the Pxx region in modulating the temperature dependence of UBQLN phase transitions.

**Figure 2.**
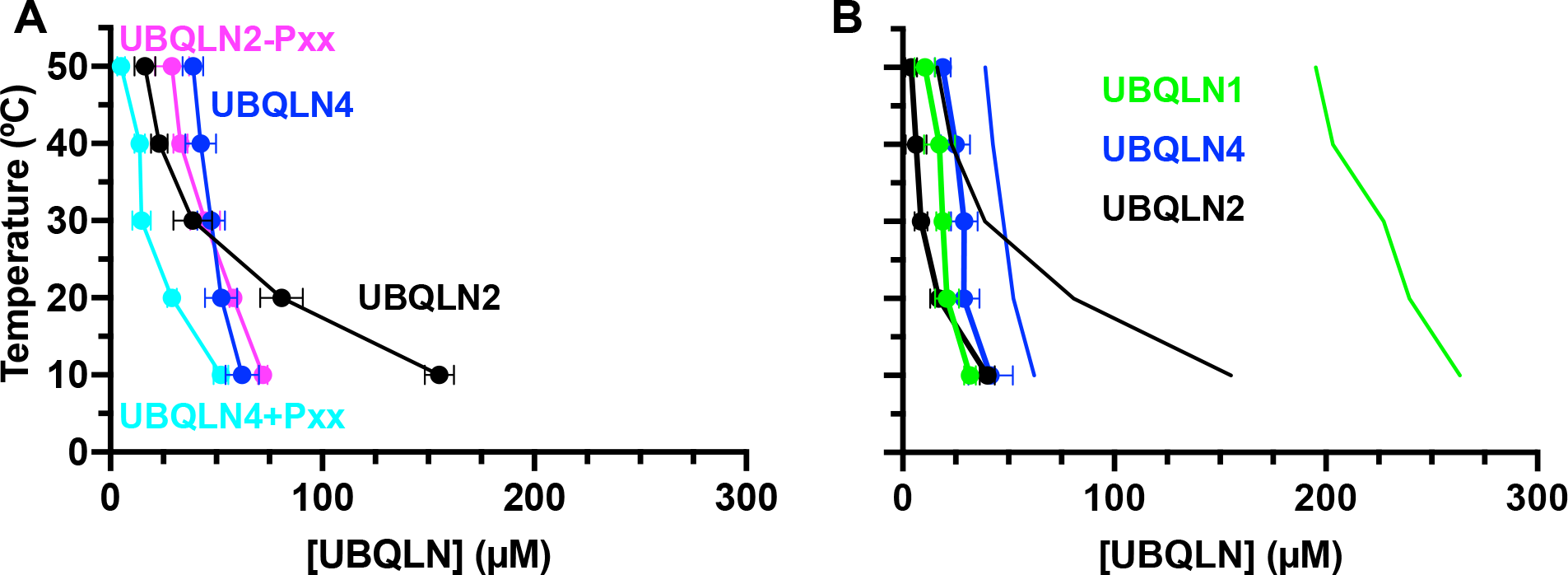
Removal of either the N-terminal disordered region or the Pxx domain alters UBQLN phase separation behaviors. Phase diagrams for (A) UBQLN2 and UBQLN4 as well as UBQLN2 without its unique Pxx region (UBQLN2-Pxx) and UBQLN4 with Pxx inserted (UBQLN4+Pxx) and (B) the three full-length UBQLNs (thin lines) and corresponding constructs without the N-terminal regions (UBQLN1 32-589, UBQLN2 33-624 and UBQLN4 12-601, filled circles, thick lines) from c_sat_ measurement as a function of temperature. Measurements were done in 20 mM NaPhosphate, 200 mM NaCl, 0.5 mM TCEP and EDTA (pH 6.8). Error bars represent SD over at least four trials from two different proteins preps.

UBQLN constructs without the N-terminal disordered regions (UBQLN1 32-589, UBQLN2 33-624 and UBQLN4 12-601) phase separated more readily than their full-length counterparts (Fig. 2B). Importantly, whereas the phase diagrams for full-length UBQLN1, UBQLN2 and UBQLN4 were highly distinguishable from each other, the phase diagrams and c_sat_ values for UBQLN1 32-589, UBQLN2 33-624 and UBQLN4 12-601 were very similar. One noticeable difference is the higher temperature dependence in the phase diagram of UBQLN2 33-624 (similar to that of UBQLN4+Pxx), further supporting the role of the Pxx region in regulating the temperature dependence of UBQLN2 phase separation. Together, these data demonstrated that the N-terminal disordered regions of UBQLN1, UBQLN2 and UBQLN4 inhibit phase separation of the three proteins and that differences in these regions contribute substantially to the differences in the phase transitions of full-length UBQLNs.

### Charged residues within the N-terminal disordered regions dictate UBQLN phase separation propensity

Upon inspecting the sequences of the UBQLN N-terminal disordered regions, we noted large differences in their charge states, especially between UBQLN1 and UBQLN2 (Fig. 3A). The N-terminal regions of UBQLN1 and UBQLN2 are highly similar, except at positions 12, 15 and 21, which are either neutral or negatively charged for UBQLN1 and positively charged or neutral for UBQLN2. These differences result in net charges at neutral pH of -4 and 0 for UBQLN1 and UBQLN2 N-terminal regions, respectively. We hypothesized that negatively charged amino acids could lead to the low phase separation propensity of UBQLN1 compared to that of UBQLN2.

**Figure 3.**
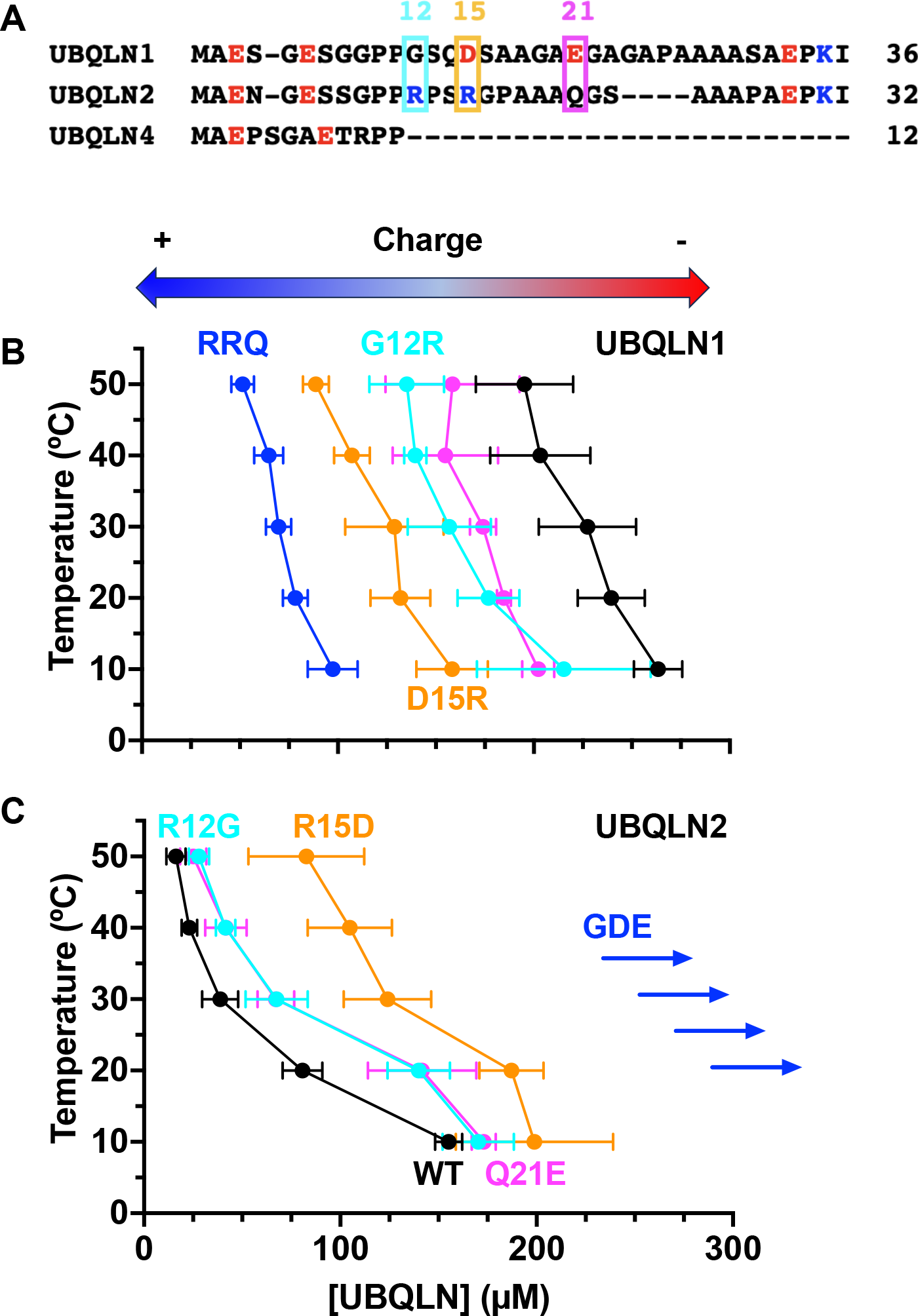
More negatively charged N-terminal regions decrease UBQLN phase separating ability. (A) Sequence alignment of the N-terminal disordered regions of UBQLN1, UBQLN2 and UBQLN4. Basic residues are in blue and acidic residues are in red. (B, C) Phase diagrams for (B) UBQLN1 and (C) UBQLN2 charge variants from c_sat_ measurements as a function of temperature. The arrows for UBQLN2 GDE variant indicate much higher c_sat_ than the range tested here. Measurements were done in 20 mM NaPhosphate, 200 mM NaCl, 0.5 mM TCEP and EDTA (pH 6.8). Error bars represent SD over at least four trials from two different proteins preps.

To determine the effect of the N-terminal ionizable residues on UBQLN1 and UBQLN2 phase separation, we generated UBQLN1 and UBQLN2 charge variants at N-terminal positions 12, 15 and 21. We subsequently obtained phase diagrams of these charge variants. Specifically for UBQLN1, we made single amino acid substitutions G12R, D15R or E21Q as well as a triple variant G12R/D15R/E21Q (RRQ). Similarly for UBQLN2, we generated single amino acid substitutions R12G, R15D or Q21E as well as the triple variant R12G/R15D/Q21E (GDE). The UBQLN1 RRQ variant mimics the UBQLN2 N-terminal sequence, whereas the UBQLN2 GDE variant mimics the UBQLN1 N-terminal sequence. Remarkably, for both UBQLN1 and UBQLN2, more negatively charged N-terminal regions led to higher c_sat_ values while neutral N-terminal regions led to lower c_sat_ values (Fig. 3B and 3C). In other words, more negative charge on the N-terminus correlates with decreased propensity to phase separate. Indeed, UBQLN2 GDE (with a net charge of -4) does not phase separate up to at least 500 µM at any temperature under the same buffer condition. Strikingly, the c_sat_ values for the UBQLN1 RRQ variant, whose N-terminal sequence mimics that of UBQLN2, are now within the same range as wild-type UBQLN2 (Fig. 3B and 3C). These data illustrate that the charge state of the UBQLN N-terminal regions is an important determinant of the propensity for UBQLNs to phase separate.

### Epitope tags alter the ability of UBQLNs to phase separate

Given the sensitivity of UBQLN phase separation to the presence or composition of the N-terminal regions, we investigated the effects of different N-terminal epitope tags, which are necessary for many studies in cells, on the propensity of UBQLNs to phase separate. We chose the commonly used FLAG, HA and Myc tags, which all carry a negative charge, and the charge-neutral ALFA tag (Götzke et al., 2019) (Fig. 4A). We added the tags on either the N-terminus of UBQLN2 or UBQLN4 and purified the constructs using our salting out protocol (see Experimental Procedures). We were able to purify all proteins except for Myc-UBQLN2, the solution of which did not turn cloudy under high NaCl and temperature conditions like the others. This result indicates that Myc-UBQLN2 does not phase separate or has a very low phase separation propensity under these conditions.

**Figure 4.**
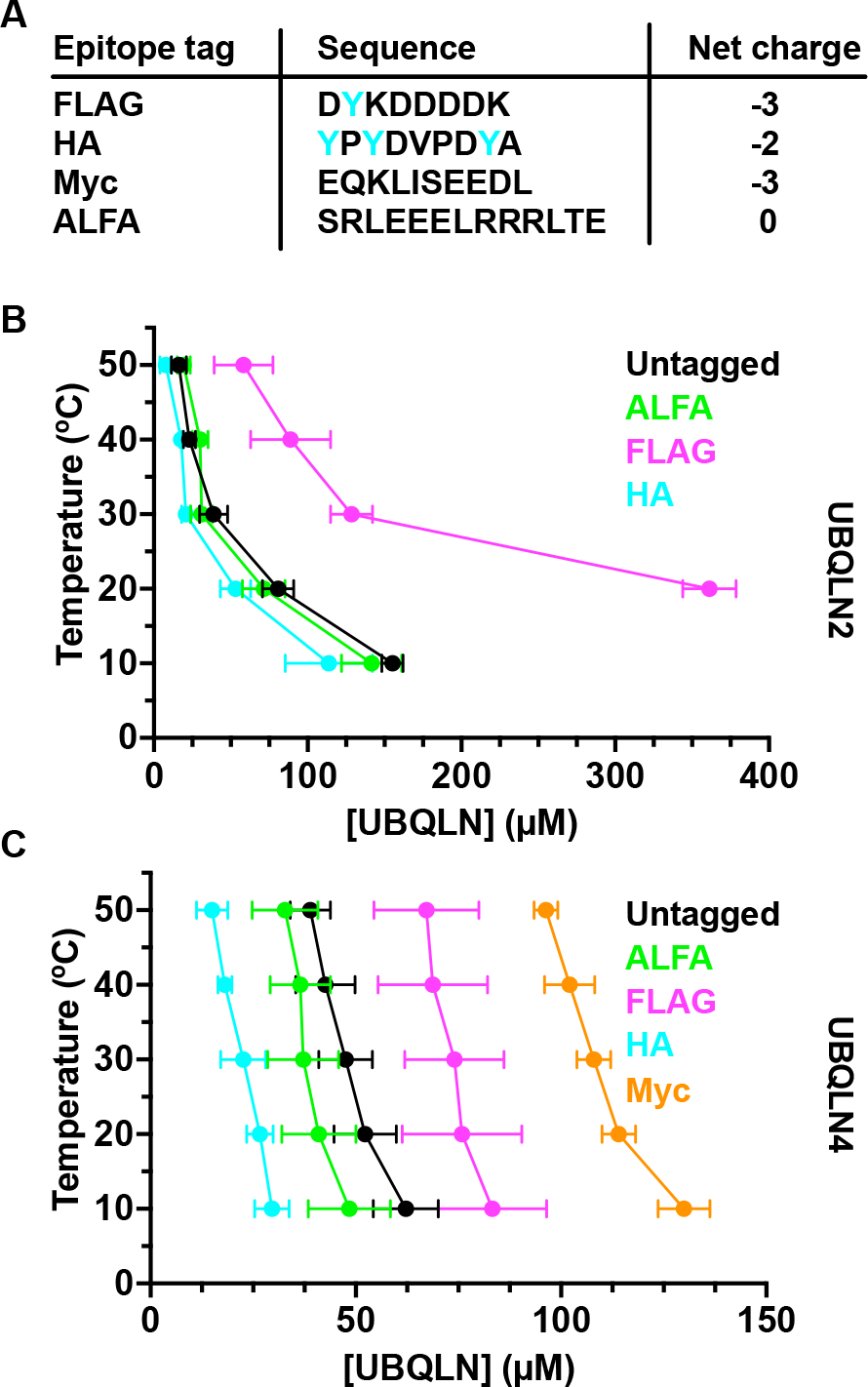
Epitope tags affect UBQLN phase separation to varying degrees. (A) Sequences and net charges of epitope tags used in this study (tyrosine residues in cyan). (B,C) Phase diagrams for (B) UBQLN2 and (C) UBQLN4 with different epitope tags from c_sat_ measurement as a function of temperature. Myc-UBQLN2 is not included as it did not phase separate under these conditions. Measurements were done in 20 mM NaPhosphate, 200 mM NaCl, 0.5 mM TCEP and EDTA (pH 6.8). Error bars represent SD over at least four trials from two different proteins preps.

Data from temperature-concentration phase diagrams revealed a wide range of effects from installing the N-terminal tags on UBQLN phase separation, and we observed the same trends for both UBQLN2 and UBQLN4 (Fig. 4B and 4C). Myc and FLAG tags inhibited UBQLN phase separation, with Myc tag to a larger extent (Myc-UBQLN2 did not phase separate, even at much higher salt and protein concentrations). ALFA and HA tags slightly and greatly enhanced phase separation, respectively. These results were inconsistent with the similarity in the negative charges of FLAG, HA and Myc tags. However, HA, FLAG and Myc tags also contain three, one and zero tyrosine residues, respectively. We previously showed that the presence of aromatic residues substantially enhances UBQLN2 phase separation (Yang et al., 2019). Our data suggest that the phase separation-promoting effects of the tyrosine residues compensate for the inhibiting effects of the overall negative charges of the HA and FLAG tags. It is likely that the tyrosines enable HA tag to enhance UBQLN phase separation the most, while the single tyrosine in FLAG tag reduces the inhibition of phase separation compared to the tyrosine-free Myc tag. Overall, the tyrosine-free, charge-neutral ALFA tag perturbs UBQLN phase separation the least.

## Discussion

*UBQLN* genes are found in all eukaryotes, with most species having one single gene (Marín, 2014). However, mammals contain multiple lineage-specific duplications, resulting in the existence of the highly homologous UBQLN1, UBQLN2 and UBQLN4, likely due to the need for new functions (Marín, 2014). How are these three proteins distinguished in the cell to carry out their specific functions, given the high sequence identity and similar domain architectures among them? One way is through tissue-specific expression. Whereas UBQLN1 is similarly expressed in all tissues, UBQLN2 is highly expressed in the nervous system (Marín, 2014). Another way is through interactions with specific binding partners and client proteins (Zheng et al., 2020). Here, we present a third possible mechanism through the distinct phase transitions of UBQLN1, UBQLN2 and UBQLN4. Aberrant phase transitions of proteins associated with neurodegeneration lead to aggregation and disease (Alberti and Hyman, 2021; Li et al., 2022; Molliex et al., 2015; Patel et al., 2015). As all three UBQLNs are involved in neurodegenerative diseases and colocalized with cellular condensates and protein aggregates, their different propensities to undergo phase transitions can contribute to their similar but distinct cellular functions and possibly disease states when dysregulated.

As expected for its uniqueness, the Pxx region of UBQLN2 plays a major role in distinguishing its phase transition from those of its close paralogs, namely by conferring a high temperature sensitivity to the UBQLN2 phase transition between 10 and 50 °C (Fig. 2A). The Pxx region, only found in mammalian UBQLN2 but not any other UBQLN paralogs or orthologs, harbors the majority of the familial ALS-linked mutations in UBQLN2 (Lin et al., 2021; Renaud et al., 2019; Zheng et al., 2020). These mutations change the phase separation propensities and material properties of UBQLN2 condensates *in vitro* and in cells (Dao et al., 2019; Riley et al., 2021; Sharkey et al., 2018). Together, these data suggest that the ability of the Pxx region to regulate the UBQLN2 phase transition is important for the functions of UBQLN2 and overall cellular health. The mechanism for how the highly temperature-dependent phase transition of UBQLN2 affects its functions is unknown. However, UBQLN2, but not UBQLN1 or UBQLN4, is associated with stress granules, particularly ones that form under heat stress (Alexander et al., 2018; Dao et al., 2018). The presence of UBQLN2 in stress granules can help regulate the material properties or disassembly of stress granules, and therefore is important for maintaining homeostasis. Another proline-rich region, the P domain of Pab1 in budding yeast, modulates the temperature dependency of the Pab1 phase transition (Riback et al., 2017). Mutations that disrupt this temperature dependency reduce organism fitness during prolonged high temperature growth. Results from studies of Pxx mutants in cells have been complicated by overexpression, which leads to the mislocalization of UBQLN2 (Riley et al., 2021), and by the addition of epitope or fluorescence tags, which, as we show in this work, can affect the phase transition of UBQLN2. Therefore, it is essential to study Pxx mutants at endogenous expression levels in cultured neurons and model organisms, to determine the biological role of the Pxx region as well as the significance of the temperature sensitivity of the UBQLN2 phase transition.

Our findings that the N-terminal disordered regions of UBQLN1, UBQLN2 and UBQLN4 inhibit and distinguish their phase transitions (Fig. 2B) were quite surprising since these regions are short and contain no obvious structural or conserved sequence features (Fig. 3A). We had previously shown that multiple UBQLN2 regions, including the oligomerization domain STI1-II, putative heat-shock protein binding domain STI1-I, and the folded UBL and UBA domains, are important for modulating its phase transitions (Dao et al., 2018; Zheng et al., 2021). These regions are referred to as stickers (Pappu et al., 2023; Rubinstein and Dobrynin, 1997). For UBQLN2, the stickers comprise mostly hydrophobic and polar residues (Dao et al., 2019, 2018). Substitutions of stickers to aromatic, positively charged or more hydrophobic residues enhance UBQLN2 phase separation, while substitutions to acidic residues substantially inhibit phase separation (Yang et al., 2019). A UBQLN1 construct with its arginine residues replaced with neutral glutamine residues is also unable to form condensates in cells (Kampmeyer et al., 2023). Consistent with these trends, UBQLN1 or UBQLN2 constructs with a negatively charged residue in the N-terminal sticker region replaced with a non-ionizable residue or a positively charged arginine exhibit higher propensities to phase separate (lower c_sat_) (Fig. 3B and 3C). Moreover, inserting sequences in front of the N-terminal sticker regions also modulates UBQLN phase transitions in a predictable manner. Specifically, negatively charged amino acids inhibit phase separation (e.g., FLAG and Myc tags, Fig. 4). However, the presence of aromatic residue tyrosine likely reduces the inhibitory effects of the negatively charged amino acids and can even lead to the promotion of phase separation in the case of the HA tag. Therefore, UBQLN phase transitions are very sensitive to the sequence of the N-terminal region; potential binding partners, mutations, or post-translational modifications such as phosphorylation (a few sites exist on the N-terminal regions of UBQLN1 and UBQLN2, Hornbeck et al., 2015) therefore likely drastically alter the localizations and functions of these proteins.

Our findings that many commonly used epitope tags alter the phase transitions of UBQLNs could have significant implications for the design of cell-based studies, especially if the underlying mechanism relies on the ability of UBQLNs (and possibly other phase-separating proteins) to form condensates and aggregates. Of the four epitope tags studied, the charge-neutral ALFA tag appears to be the most ideal for placing on the N-terminus of UBQLNs as it minimally affects the phase transitions of UBQLN2 and UBQLN4 (Fig. 4). Moreover, the ALFA tag is recognized by the easily accessible and highly specific nanobody NbALFA, which can be expressed and purified from bacterial cells and modified with fluorescent proteins or organic dyes for detection in cells (Götzke et al., 2019). However, the best epitope tag to use is most likely protein-dependent. The ALFA tag, being helical in nature, may not be ideal for proteins whose phase separation is dependent on higher helical propensities (Chiu et al., 2022; Conicella et al., 2020). Another factor to consider is the location of a tag. Our data suggest that the N-termini of UBQLNs are not good locations for epitope tags as these N-terminal regions are stickers that modulate UBQLN phase transitions. Phase-separating proteins also contain spacers, non-interacting sequences that connect stickers (Choi et al., 2020). Changes to spacer regions within UBQLN2 are much less disruptive to phase transitions (Yang et al., 2019), suggesting that spacer regions might be better for the placement of epitope tags. However, care must be taken to avoid disrupting known binding sites of interacting partners. Ideally, cell work should be preceded by an *in vitro* test to ensure that the phase transition of the target protein remains largely unchanged with an epitope tag placed in a certain location.

## Conclusions

In this work, we demonstrated that full-length UBQLN1, UBQLN2 and UBQLN4 are all capable of phase separation, but to very different extents. Furthermore, we identified the previously uncharacterized N-terminal disordered region of each protein and the Pxx domain of UBQLN2 as stickers that not only modulate but also distinguish the phase transitions of these three proteins. Without these regions, UBQLN1, UBQLN2 and UBQLN4 exhibit similar phase transitions with c_sat_ values in the low µM range. Our study suggests that even short N-terminal (or C-terminal) disordered regions must be carefully examined when determining the driving forces of phase separation. Phase transitions can be very sensitive to single amino acid substitutions or addition of short but specific sequences.

## Experimental Procedures

### Subcloning, Protein Expression, and Purification

The genes encoding UBQLN1, UBQLN2 and UBQLN4 constructs were codon-optimized, synthesized, and cloned into pET24b (Novagen) by GenScript (NJ, USA). All variant UBQLN constructs were made using Phusion Site-Directed Mutagenesis Kit (Thermo Scientific). UBQLN constructs were grown in NiCo21 cells in Luria-Bertani (LB) broth with 50 mg/L kanamycin to OD_600_ of 0.6, induced with 0.5 mM IPTG and expressed overnight at 37°C (except for UBQLN1, UBQLN1 G12R, UBQLN1 D15R and UBQLN1 E21Q, which were expressed at 20 °C).

Cells were pelleted, frozen, and resuspended in 20 mL (for every 2 L of cells) of 50 mM Tris buffer pH 8.0 with 1 mM PMSF, 2 mM MgCl_2_, 1 mg/ml lysozyme and 10 U of Pierce universal nuclease. The lysate was cleared by centrifugation at 20 000 *g for 30 min. All constructs, except for UBQLN2 GDE and Myc-UBQLN2, were purified via a “salting out” process, where NaCl was added to the lysate to a final concentration of 0.5 -2 M. The resulting UBQLN dense phase was pelleted and then resuspended in cold 20 mM sodium phosphate buffer pH 6.8, with 0.5 mM EDTA & 0.02 % NaN_3_. Solutions of UBQLN2 and UBQLN4 constructs as well as UBQLN1 32-589 were further cleaned up through another round of salting out and solubilization of pelleted dense phase into cold 20 mM sodium phosphate buffer pH 6.8, with 0.5 mM EDTA & 0.02 % NaN_3_ or 20 mM Hepes buffer pH 7 (UBQLN1 32-589, UBQLN2 33-624, UBQLN4 12-601). Solutions of UBQLN1, UBQLN1 G12R, UBQLN1 D15R and UBQLN1 E21Q were subjected to size exclusion chromatography over an ENrich™ SEC 650 10 x 300 column (BioRad) in 20 mM sodium phosphate buffer pH 6.8, with 0.5 mM EDTA & 0.02 % NaN_3_. Proteins solutions were concentrated to at least 600 µM with Vivaspin centrifugal concentrators (Sartorius) and stored at -80 °C.

For UBQLN2 GDE and Myc-UBQLN2, lysate was loaded onto an anion exchange column (Cytiva) and eluted with a gradient (Buffer A: 20 mM Hepes, pH 7.5, Buffer B: 20 mM Hepes, pH 7.5, 1 M NaCl). Fractions collected were diluted into 50 mM ammonium acetate, pH 4.5 until solution turned cloudy. Pellet was washed and redissolved into 20 mM sodium phosphate buffer pH 6.8, with 0.5 mM EDTA & 0.02 % NaN_3_.

### Bright-field Imaging of Phase Separation

Samples were prepared to contain 200 μM UBQLNs in 20 mM NaPhosphate, 200 mM NaCl, 0.1 mM TCEP, and 0.5 mM EDTA (pH 6.8). Samples were added to Eisco Labs Microscope Slides, with Single Concavity, and covered with MatTek coverslips that had been coated with 5% bovine serum albumin (BSA) to minimize changes due to surface interactions, and incubated coverslip-side down at 30 °C for 5 min. Phase separation was imaged on an ONI Nanoimager (Oxford Nanoimaging Ltd, Oxford, UK) equipped with a Hamamatsu sCMOS ORCA flash 4.0 V3 camera using an Olympus 100×/1.4 N.A. objective. Images were prepared using Fiji (Schindelin et al., 2012) and FigureJ plugin.

### Phase Diagram Measurements

We quantified the saturation concentration (c_sat_) values for the different UBQLN constructs using SDS-PAGE gels. 15 µL of 2X stocks of UBQLN constructs (280-400 µM UBQLN1 32-589, UBQLN2 33-624, UBQLN4 12-601; 600 µM for UBQLN1, UBQLN1 G12R, D15R, E21Q, UBQLN2 R15D; 725-960 µM for FLAG-UBQLN2; 400 µM for the rest of the constructs) were mixed with 15 µL of 2X salt solution (20 mM NaPhosphate (pH 6.8), 0.5 mM TCEP, 0.5 mM EDTA, 400 mM NaCl) and incubated for 10 minutes in temperature-equilibrated centrifuges at 10, 20, 30, 40 and 50 °C. Samples were then centrifuged for 5 minutes at 21000 x g. 1 µL of the supernatant of each sample was immediately mixed with 20 µL of 2X SDS-PAGE dye. 4 µL of each sample were loaded onto 10% polyacrylamide gels, imaged using the BioRad Gel Doc EZ Imager, and band volumes were determined with BioRad Image Lab software. For each gel, samples for a standard curve containing the solution concentration for a particular experiment (c_sol_), c_sol_/2, c_sol_/4, c_sol_ /8, c_sol_/16 µM of protein were also loaded, analyzed and used to calculate the c_sat_ values of the proteins. The measurements were carried out using at least two different protein stocks with two trials for each.

### Western Blotting

A total of 250 ng of purified proteins as labelled above each lane were loaded on TGX stain-free gels (BioRad) and run at constant 150V. Blotting was done by standard mini-TGX protocol (2.5 A, 25 V, 3 minutes) using a Trans-blot Turbo Transfer System (BioRad). Membranes were incubated with UBQLN2 primary antibodies (1:5000 diluted) in Intercept® (PBS) Blocking Buffer containing 0.2% Tween20 at 4°C overnight. Primary antibodies include UBQLN2 rabbit mAb (Cell Signaling Technology 85509), rabbit UBQLN2 pAb (Proteintech 23449-1-AP) or UBQLN2 mouse mAb (Novus Biologicals NBP2-25164). After three washes, membranes were incubated with secondary antibodies (LI-COR Biosciences, IRDye® 800CW Goat anti-Mouse IgG Secondary Antibody-925-32210 and IRDye® 680RD Goat anti-Rabbit IgG Secondary Antibody-926-68071) diluted 1:5000 in blocking buffer containing 0.2% Tween 20 and 0.02% SDS at room temperature for two hours. After three additional washes, bands were visualized with a ChemiDoc MP Imaging system (BioRad).

### SEC-MALS-SAXS Experiments

SAXS was performed at BioCAT (beamline 18ID at the Advanced Photon Source, Chicago) with in-line size exclusion chromatography (SEC-SAXS) to separate sample from aggregates and other contaminants thus ensuring optimal sample quality and multiangle light scattering (MALS), dynamic light scattering (DLS) and refractive index measurement (RI) for additional biophysical characterization (SEC-MALS-SAXS). Sample was loaded onto a Superdex 200 Increase 10/300 GL column (Cytiva) run by 1260 Infinity II HPLC (Agilent Technologies) at 0.6 ml/min. The flow passed through (in order) the Agilent UV detector, a MALS detector and a DLS detector (DAWN Helios II, Wyatt Technologies), and an RI detector (Optilab T-rEX, Wyatt). The flow then went through the SAXS flow cell. The flow cell consists of a 1.0 mm ID quartz capillary with ∼20 µm walls. A coflowing buffer sheath is used to separate the sample from the capillary walls, helping prevent radiation damage (Kirby et al., 2016). Scattering intensity was recorded using a Pilatus3 X 1 M (dectris) detector which was placed 3.6 m from the sample giving access to a q-range of 0.003 Å^-1^ to 0.35 Å^-1^. 0.5 s exposure was acquired every 2 s during elution, and data were reduced using BioXTAS RAW 2.1.1 (Hopkins et al., 2017). Buffer blanks were created by averaging regions flanking the elution peak (see Fig S4) and subtracted from exposure selected from the elution peak to create the I(*q*) vs. *q* curves used for subsequent analysis. Molecular weights and hydrodynamic radii were calculated from the MALS and DLS data respectively using ASTRA 7 software (Wyatt). Additionally, R_g_ and I(0) values were obtained using the entire q-range of the data by calculating the distance distribution functions, P(*r*) vs. r, using GNOM (Svergun, 1992). All SEC-MALS-SAXS parameters for data collection and analysis can be found in Table S2.

## Data availability

SAXS data are deposited in SASBDB in two project files: https://www.sasbdb.org/project/2151/ (UBQLN2-PXX, UBQLN2 ΔSTI1-II) and https://www.sasbdb.org/project/2068/ (UBQLN1, UBQLN2, UBQLN4). All other data are available from corresponding author upon request.

## Supporting information

This article contains supporting information.

## Acknowledgements

We acknowledge support and assistance from Dr. Jesse Hopkins and Dr. Maxwell Watkins on collecting SEC-MALS-SAXS data at APS. We acknowledge Drs. Yiran Yang and Tongyin Zheng for initial work on UBQLN1 and UBQLN4. We thank Dr. Dan Kraut for critical reading and feedback on the manuscript.

## Funding and additional information

This work was supported by NSF CAREER award MCB-1750462 (all protein purifications, phase diagrams) and NIH R01GM136946 (western blots, SEC-MALS-SAXS) to C.A.C.. This research used resources of the Advanced Photon Source, a U.S. Department of Energy (DOE) Office of Science User Facility operated for the DOE Office of Science by Argonne National Laboratory under Contract No. DE-AC02-06CH11357. This project was supported by grant P30 GM138395 from the National Institute of General Medical Sciences of the National Institutes of Health. Use of the Pilatus 3 1 M detector was provided by grant 1S10OD018090-01 from NIGMS. The content is solely the responsibility of the authors and does not necessarily represent the official views of the National Institutes of Health.

