## Supporting Information for "Short N-terminal disordered regions and the proline-rich domain are major regulators of phase transitions for full-length UBQLN1, UBQLN2 and UBQLN4"

Contents:

Figures S1 – S6
Tables S1 – S3
References

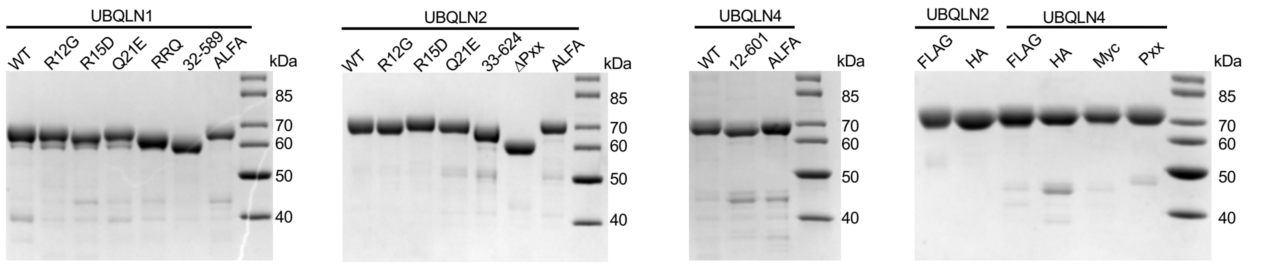


**Figure S1**. Representative SDS-PAGE gel images of the three purified UBQLNs and variants used in this study. Molecular weight markers are shown in the last lane of each gel.

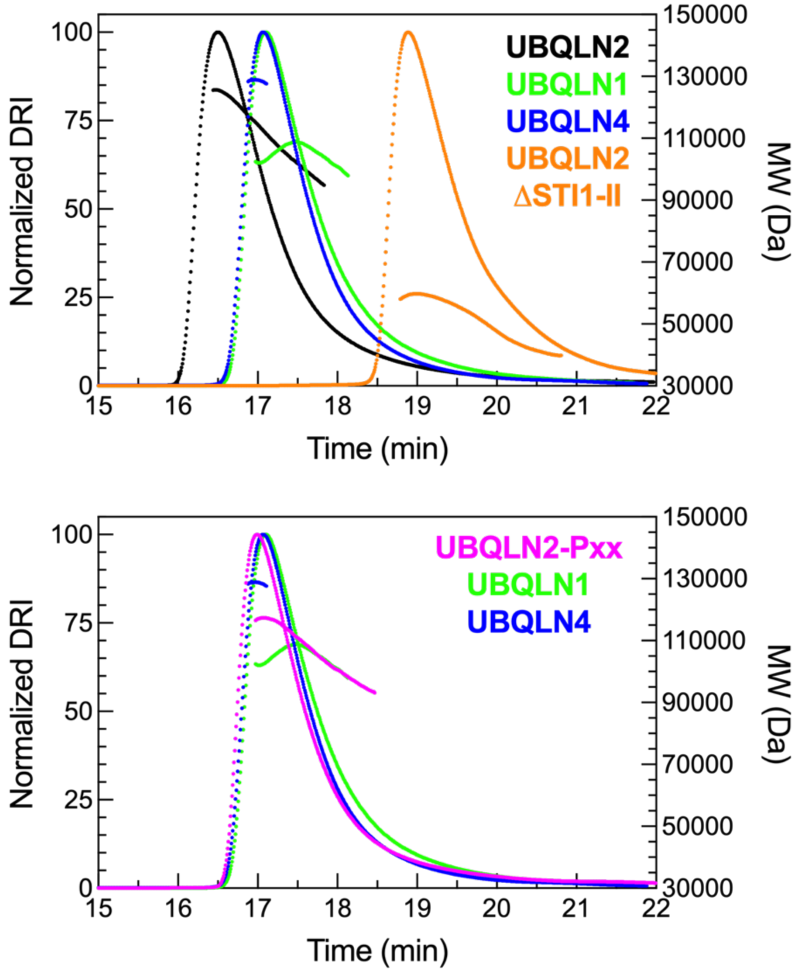


**Figure S2.** SEC-MALS profiles for UBQLN1, UBQLN2, and UBQLN4, and two deletion constructs of UBQLN2. The MALS-derived molecular weights for full-length UBQLN1, UBQLN2, and UBQLN4 are consistent with these proteins existing in the dimeric state. For comparison, we include UBQLN2-ΔSTI1-II (residues 379-462 removed) as it is monomeric. These data are in agreement with our previous report (Dao et al., 2018) that showed UBQLN2 was monomeric when residues 379-486 were removed. Removal of the proline-rich region (UBQLN2-Pxx) did not alter the oligomerization state of UBQLN2.

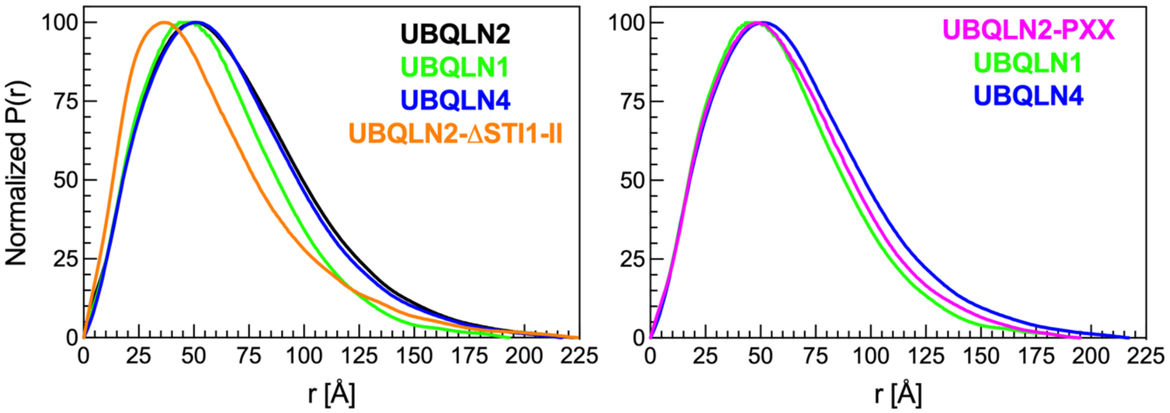


**Fig. S3.** Normalized P(r) profiles from SAXS experiments (see Fig. S4) of various UBQLN constructs.

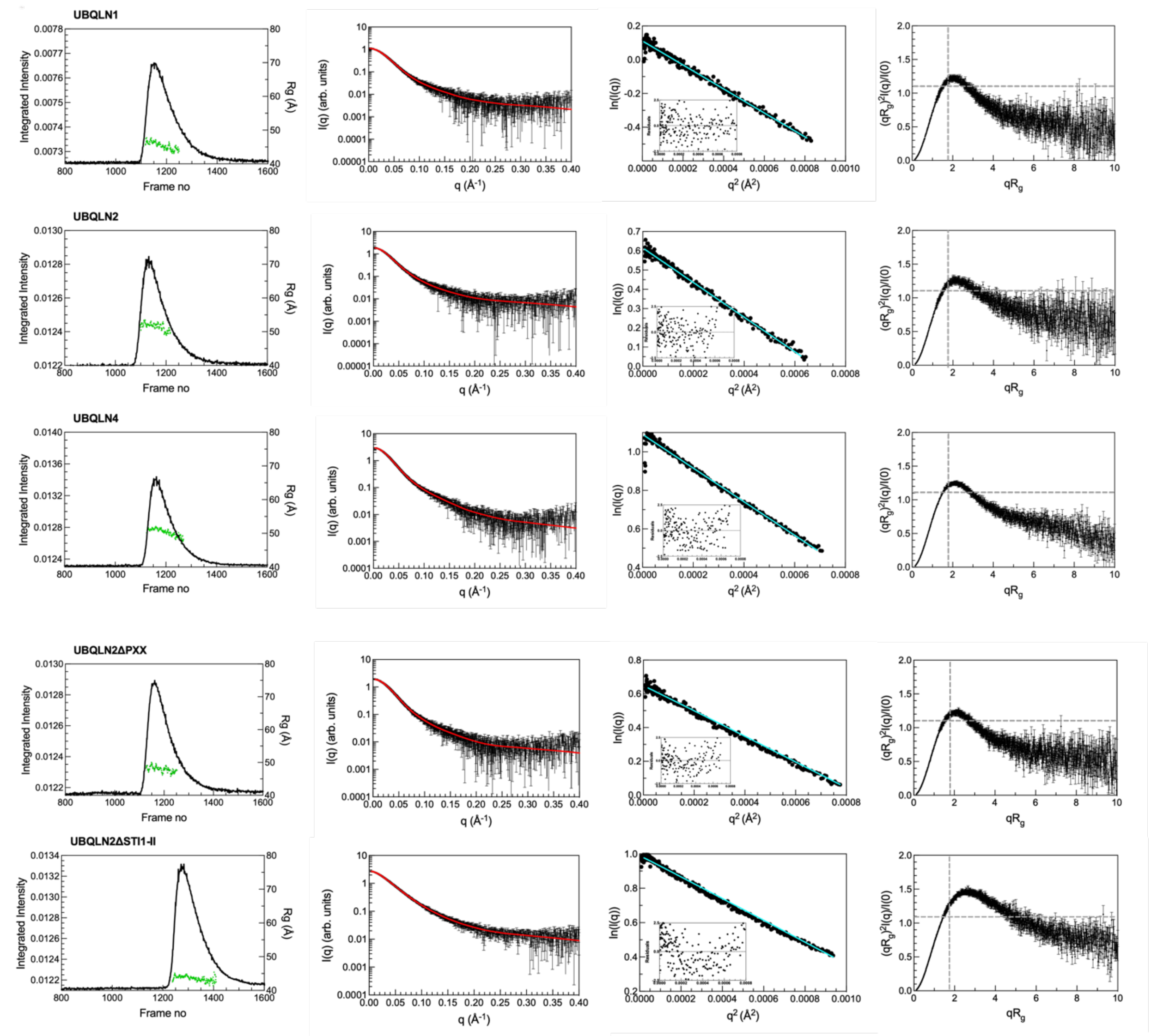


**Figure S4.** Solution scattering data for full-length UBQLNs. (Left) SEC-SAXS profiles for UBQLN1, UBQLN2, UBQLN4, UBQLN2-PXX, and UBQLN2 ΔSTI1-II. (Middle left) I(q) vs. q scattering curves determined from frames 1133-1159 for UBQLN1, frames 1104-1129 for UBQLN2, frames 1141-1181 for UBQLN4, frames 1150-1174 for UBQLN2-Pxx, and frames 1259-1285 for UBQLN2 ΔSTI1-II. Red line in the I(q) profile indicates fit from P(r) analysis (see Figure S3). (Middle right) Guinier plot with linear fit (cyan) of ln(I(q)) vs. $q^2$, while inset shows residuals of fit. (Right) Dimensionless Kratky plots include dashed lines to indicate where a globular protein would peak.

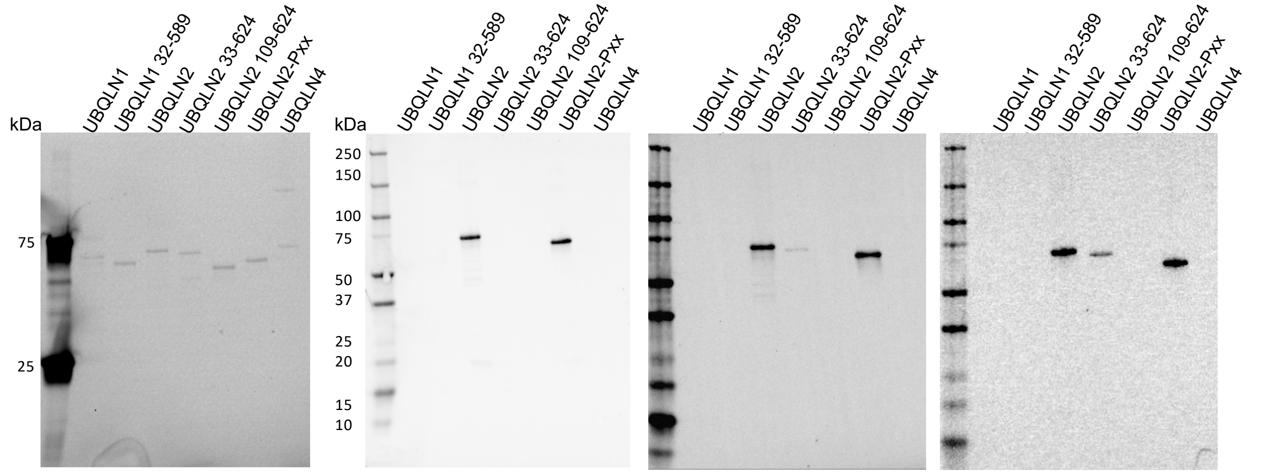


Representative TGX stain-free gel prior to transfer

CST UBQLN2 (D7R2Z) Rabbit mAb #85509 Dilution 1:5000

Novus UBQLN2 (6H9) Mouse mAb NBP2-25164SS Dilution 1:5000

Proteintech UBQLN2 Rabbit pAb 23449-1-AP Dilution 1:5000

**Figure S5.** Western blots showing that the epitope for three distinct UBQLN2-specific antibodies is mainly located within the first 32 residues of UBQLN2. Constructs are described in Table S3.

```
UBQLN2      MAEN-GESSGPPRPSRGPAAAQGSAAAPAEPKIIKVTVKTPKEKEEFAVPENSSVQQFKEAISKRFKSQTDQLVLIFAGKILKDQDTLIQHGIHDGLTVH    99
UBQLN2-Pxx  MAEN-GESSGPPRPSRGPAAAQGSAAAPAEPKIIKVTVKTPKEKEEFAVPENSSVQQFKEAISKRFKSQTDQLVLIFAGKILKDQDTLIQHGIHDGLTVH    99
UBQLN4      MAEPSGAETRPP---------------------IRVTVKTPKDKEEIVICDRASVKEFKEEISRRFKAQQDQLVLIFAGKILKDGDTLNQHGIKDGLTVH    79
UBQLN4+Pxx  MAEPSGAETRPP---------------------IRVTVKTPKDKEEIVICDRASVKEFKEEISRRFKAQQDQLVLIFAGKILKDGDTLNQHGIKDGLTVH    79
            ***  * .: **                     *:*******:***:.: :.:**::*** **:***:* ************** *** ****:******

UBQLN2      LVIKSQNRPQGQS---------TQPSNAAGTNTTSASTPRSNSTPISTNSNP-------------------------FGLGSLGGLAGLSSLGLSSTNFS   165
UBQLN2-Pxx  LVIKSQNRPQGQS---------TQPSNAAGTNTTSASTPRSNSTPISTNSNP-------------------------FGLGSLGGLAGLSSLGLSSTNFS   165
UBQLN4      LVIKTPQKAQDPAAATASSPSTPDPASAPSTTPASPATPAQPSTSGSASSDAGSGSRRSSGGGPSPGAGEGSPSATASILSGFGGILGLGSLGLGSANFM   179
UBQLN4+Pxx  LVIKTPQKAQDPAAATASSPSTPDPASAPSTTPASPATPAQPSTSGSASSDAGSGSRRSSGGGPSPGAGEGSPSATASILSGFGGILGLGSLGLGSANFM   179
            ****: ::.*. :         .:*:.*..*..:*.:** . **. *:.*:.                           *..:**: **.****.*:**

UBQLN2      ELQSQMQQQLMASPEMMIQIMENPFVQSMLSNPDLMRQLIMANPQMQQLIQRNPEISHLLNNPDIMRQTLEIARNPAMMQEMMRNQDLALSNLESIPGGY   265
UBQLN2-Pxx  ELQSQMQQQLMASPEMMIQIMENPFVQSMLSNPDLMRQLIMANPQMQQLIQRNPEISHLLNNPDIMRQTLEIARNPAMMQEMMRNQDLALSNLESIPGGY   265
UBQLN4      ELQQQMQRQLMSNPEMLSQIMENPLVQDMMSNPDLMRHMIMANPQMQQLMERNPEISHMLNNPELMRQTMELARNPAMMQEMMRNQDRALSNLESIPGGY   279
UBQLN4+Pxx  ELQQQMQRQLMSNPEMLSQIMENPLVQDMMSNPDLMRHMIMANPQMQQLMERNPEISHMLNNPELMRQTMELARNPAMMQEMMRNQDRALSNLESIPGGY   279
            ***.***:***:.***: ******:**.*:*******::**********::*******:****::****:*:*************** ************

UBQLN2      NALRRMYTDIQEPMLNAAQEQFGGNPFASVGSSSSSGEGTQPSRTENRDPLPNPWAPPPATQSSATTSTTTSTGSGSGNSSSNA-TGNTVAAANYVASIF   364
UBQLN2-Pxx  NALRRMYTDIQEPMLNAAQEQFGGNPFASVGSSSSSGEGTQPSRTENRDPLPNPWAPPPATQSSATTSTTTSTGSGSGNSSSNA-TGNTVAAANYVASIF   364
UBQLN4      NALRRMYTDIQEPMFSAAREQFGNNPFSSLAGNSDS-SSSQPLRTENREPLPNPWSPSPPTSQAPGSGGEGTGGSGTSQVHPTVSNPFGINAASLGSGMF   378
UBQLN4+Pxx  NALRRMYTDIQEPMFSAAREQFGNNPFSSLAGNSDS-SSSQPLRTENREPLPNPWSPSPPTSQAPGSGGEGTGGSGTSQVHPTVSNPFGINAASLGSGMF   378
            **************:.**:****.***:*:...*.* ..:** *****:******:*.*.*..:. :.   : ***:.:  ... .   : **.  :.:*

UBQLN2      STPGMQSLLQQITENPQLIQNMLSAPYMRSMMQSLSQNPDLAAQMMLNSPLFTANPQLQEQMRPQLPAFLQQMQNPDTLSAMSNPRAMQALMQIQQGLQT   464
UBQLN2-Pxx  STPGMQSLLQQITENPQLIQNMLSAPYMRSMMQSLSQNPDLAAQMMLNSPLFTANPQLQEQMRPQLPAFLQQMQNPDTLSAMSNPRAMQALMQIQQGLQT   464
UBQLN4      NSPEMQALLQQISENPQLMQNVISAPYMRSMMQTLAQNPDFAAQMMVNVPLFAGNPQLQEQLRLQLPVFLQQMQNPESLSILTNPRAMQALLQIQQGLQT   478
UBQLN4+Pxx  NSPEMQALLQQISENPQLMQNVISAPYMRSMMQTLAQNPDFAAQMMVNVPLFAGNPQLQEQLRLQLPVFLQQMQNPESLSILTNPRAMQALLQIQQGLQT   478
            .:* **:*****:*****:**::**********:*:****:*****:* ***:.*******:* ***.********::** ::********:********

UBQLN2      LATEAPGLIPSFTPGVGVGVLGTAIGPVGPVTPIGPIGPIVPFTPIGPIGPIGPTGPAAPPGSTGSGGPTGPTVSSAAPSETTSPTSESGPNQQFIQQMV   564
UBQLN2-Pxx  LATEAPGLIPSFTPGVGVGVLG----------------------------------------------------SSAAPSETTSPTSESGPNQQFIQQMV   512
UBQLN4      LQTEAPGLVPSLG------SFGISRTP------------------------------APSAGSNAGSTPEAPTSSPATP-ATSSPTGASSAQQQLMQQMI   541
UBQLN4+Pxx  LQTEAPGLVPSLG------SFGISRTPVGPVTPIGPIGPIVPFTPIGPIGPIGPTGPAAPPGSNAGSTPEAPTSSPATP-ATSSPTGASSAQQQLMQQMI   571
            * ******:**:        :*                                                    *.*:*  *:***. *..:**::***:

UBQLN2      QALAGANAPQLPNPEVRFQQQLEQLNAMGFLNREANLQALIATGGDINAAIERLLGSQPS                                           624
UBQLN2-Pxx  QALAGANAPQLPNPEVRFQQQLEQLNAMGFLNREANLQALIATGGDINAAIERLLGSQPS                                           572
UBQLN4      QLLAGSGNSQVQTPEVRFQQQLEQLNSMGFINREANLQALIATGGDINAAIERLLGSQLS                                           601
UBQLN4+Pxx  QLLAGSGNSQVQTPEVRFQQQLEQLNSMGFINREANLQALIATGGDINAAIERLLGSQLS                                           631
            * ***:. .*: .*************:***:*************************** *
```

**Figure S6.** Sequence alignment of UBQLN2, UBQLN2 with the Pxx region (cyan) removed (UBQLN2-Pxx), UBQLN4 and UBQLN4 with a segment (yellow) of the Pxx region added (UBQLN4+Pxx).

**Table S1.** Structural parameters of UBQLN variants from SAXS data analysis. Indicated in parentheses are the methods/software used for $R_g$ analysis. [a] $R_g$ and errors were determined from the linear fit of ln(I(q)) vs. $q^2$ and [b] $R_g$ and errors were determined from choosing multiple values of $D_{max}$. Data collected in 20 mM sodium phosphate buffer pH 6.8 with 0.5 mM EDTA and 0.02 % $NaN_3$.

| Protein | $R_g$ (Å)[a] (Guinier) [a] | $R_g$ (Å)[b] (GNOM) [b] | $D_{max}$ (Å) (GNOM) |
|---|---|---|---|
| UBQLN1 | 46.16 ± 0.12 | 48.07 ± 0.20 | 193 |
| UBQLN2 | 51.92 ± 0.14 | 54.70 ± 0.26 | 225 |
| UBQLN4 | 51.16 ± 0.08 | 53.75 ± 0.16 | 217 |
| UBQLN2 ΔSTI1-II | 43.97 ± 0.07 | 48.81 ± 0.24 | 224 |
| UBQLN2-PXX | 48.16 ± 0.11 | 50.18 ± 0.18 | 194 |

**Table S2.** SAXS experimental parameters for all UBQLN variants. See attached Excel XLSX file.

**Table S3.** List of protein constructs used in this study.

| Construct | Residues |
|---|---|
| UBQLN1 (full-length) | 1-589 |
| UBQLN1 32-589 | 32-589 |
| UBQLN1 G12R | 1-589 with G12R |
| UBQLN1 D15R | 1-589 with D15R |
| UBQLN1 E21Q | 1-589 with E21Q |
| UBQLN1 RRQ | 1-589 with G12R, D15R, and E21Q |
| UBQLN2 (full-length) | 1-624 |
| UBQLN2 33-624 | 33-624 |
| UBQLN2 109-624 | 109-624 |
| UBQLN2-ΔSTI1-II | 1-378, 463-624 |
| UBQLN2-Pxx | 1-486, 539-624 |
| UBQLN2 R12G | 1-624 with R12G |
| UBQLN2 R15D | 1-624 with R15D |
| UBQLN2 Q21E | 1-624 with Q21E |
| UBQLN2 GDE | 1-624 with R12G, R15D, and Q21E |
| ALFA-UBQLN2 | ALFA + 1-624 |
| FLAG-UBQLN2 | FLAG + 1-624 |
| HA-UBQLN2 | HA + 1-624 |
| Myc-UBQLN2 | Myc + 1-624 |
| UBQLN4 (full-length) | 1-601 |
| UBQLN4 12-601 | 12-601 |
| ALFA-UBQLN4 | ALFA + 1-601 |
| FLAG-UBQLN4 | FLAG + 1-601 |
| HA-UBQLN4 | HA + 1-601 |
| Myc-UBQLN4 | Myc + 1-601 |
